# Chronic Exposure to Nanoplastics Alters Stem Cell Type-Specific Mechanisms, Promoting Cancer Development

**DOI:** 10.64898/2026.01.21.700811

**Authors:** Irene Barguilla, Laura Billon, Kevin Geistlich, Javier Gutiérrez, Raquel Egea, Laura Rubio, Sandrine Jeanpierre, Alba Hernández, Boris Guyot, Véronique Maguer-Satta

## Abstract

Increasing levels of nanoplastics (NPLs) in the environment raise concerns about their effects on human health. We investigated the impact of the most prevalent NPLs, namely polyethylene terephthalate (PET) and polystyrene (PS), on stem cells (SCs), which persist for decades, support tissue function, and are often implicated in cancer development. Long-term exposure to both NPLs similarly affected mammary SC features with an enhanced self-renewal and altered 3D organization, without impairing differentiation capacity. Moreover, both NPLs significantly increased invasiveness and anchorage-independent growth, albeit molecular profiling revealed distinct signatures and mechanisms, indicating a shift towards a more aggressive phenotype. NPLs also synergized with BMP2 signaling, known to be disrupted by pollutants. These findings highlight how NPLs may contribute to early pre-neoplastic changes through distinct and cooperative mechanisms.

## Introduction

Microplastics (MPLs) and nanoplastics (NPLs) have drawn considerable attention as potential threats to human health. These small plastic particles (< 5 µm for microplastics; < 1 µm for nanoplastics) are widespread in the environment leading to the continuous exposure of populations by inhalation, ingestion or dermal contact (*1*). As a consequence, MNPLs can be internalized in primary organs, translocate through physiological barriers, reach the bloodstream and potentially bioaccumulate in secondary organs. Various types of MNPLs have already been detected in human tissues and biological fluids, including the colon, lungs, liver, blood, and milk (*2*). An increasing number of studies report that NPLs can disrupt cellular processes and induce oxidative stress, inflammation, and genotoxicity (*3, 4*). Sustained over time, these effects raise concerns about physiological disruptions and potential carcinogenicity (*5*). However, a critical gap remains in our understanding of the long-term effects of NPLs, hindering accurate risk assessment (*6*). To bridge this gap, it is essential to (i) implement methodologies that replicate chronic exposure, better reflecting real-world conditions compared to the short-term approaches commonly used; (ii) use test materials representative of real-life NPLs, as their diverse formulations and physicochemical properties (*i*.*e*., surface texture, chemical additives, degradation levels) could influence their bioavailability and effects ; and (iii) generate informative data for instance on the most important hazards they present, such as carcinogenicity, or on relevant target organs (Science for Environment Policy (2023) *Nanoplastics: state of knowledge and environmental and human health impacts*. Future Brief 27. Brief produced for the European Commission DG Environment by the Science Communication Unit, UWE Bristol. https://ec.europa.eu/science-environment-policy).

In this context, stem cells (SCs) could be an interesting model to study the long-term effects of exposure to NPLs. Indeed, these cells persist in the organism for decades, sustain all organ homeostasis and are often associated with early stages of the carcinogenic process (*7-10*). The detection of NPLs in milk (*11, 12*) indicates that the breast is a secondary organ that could be seriously affected by NPLs, due to its known sensitivity to the endocrine disrupting-chemicals present in some plastic polymers and NPLs (*13*). Our team is specialized in human mammary stem cells (MaSCs) and their involvement in cancer development and progression, and we therefore addressed this knowledge gap by developing an innovative and relevant *in vitro* model through the chronic exposure of MaSCs to NPLs for up to 20 weeks. Using a comprehensive panel of validated assays (*14*), we characterized changes in stemness, molecular alterations, and transformation phenotypes induced by long-term NPL exposure. We also investigated how NPLs may cooperate with soluble bone morphogenetic protein 2 (BMP2) signaling in MaSC transformation, as BMP2, present in the breast microenvironment, regulates healthy MaSC function but is also implicated in their transformation (*7, 15*). Of note, we previously demonstrated that BMP signaling can be disrupted by long-term exposure to environmental pollutants, such as endocrine disruptors like bisphenol A or BenzoApyrene (*16*). This study is the first to compare the long-term effects of pristine polystyrene nanobeads (PS) and real-life polyethylene terephthalate nanoplastics (PET) on human SCs.

## Materials and Methods

### PS and PET nanoplastic obtention and characterization

PS was purchased as a stable dispersion of 50 nm nanoplastic particles (PP-008-10) from Spherotech. PET nanoplastic was obtained in-house by milling commercial PET water bottles. Briefly, bottles were sanded with a rotary diamond burr and sieved through a 0.22-mm mesh. Four grams of the resulting particles were added to 40 mL of 50 °C pre-warmed 90 % trifluoroacetic acid with constant stirring for a 2 h and then kept under constant agitation overnight. The pellet was resuspended in 400 mL of 0.50 % sodium dodecyl sulphate, sonicated, and allowed to settle in 250-mL graduated cylinders for 1 h. Stock suspensions of 180 nm in size at 5 mg/mL were prepared for use in cell culture experiments. Fluorescently labelled versions of PS and PET nanoplastic were also used for internalization assessment. Yellow-fluorescent PS nanoplastic (FP-00552-2) comparable to the non-labelled PS was purchased from Spherotech. Alternatively, PET nanoplastic particles were labelled with iDye Poly Pink (Rupert, Gibbon & Spider, Inc.). Briefly, 0.01 g of iDye Poly Pink was incubated together with 1 mL of the stock PET suspension at 70°C for 2 h. After cooling, the suspension was washed by adding it to 9 mL of Mili-Q water in an Amicon Ultra-15 Centrifugal Filter Unit (Merck) and centrifuged at 4,000 rpm for 15 min. After two additional washing steps, the particles were collected and suspended in a final volume of 1 mL of Mili-Q water.

### Cell culture conditions

MCF10A cells were purchased from ATCC and cultured in phenol red-free Dulbeco’s modified Eagle’s medium (DMEM)/F-12 (Thermo Fischer Scientific) supplemented with 1% penicillin/streptomycin, 5% horse serum, 10 µg/mL insulin, 0.5 µg/mL hydrocortisone, 100 ng/mL cholera toxin, and 20 ng/mL EGF (Sigma-Aldrich) in a humidified atmosphere of 5% CO_2_ and 95% air at 37ºC.

### Nanoplastic internalization

30,000 cells were seeded in top of sterile coverslips placed on 12-well culture plates. The cells were exposed to 100 µg/mL fluorescent PS and PET for 24, 48 and 72 h. Then, cells were washed twice with cold PBS 1X, fixed with PFA 4% for 20 min and permeabilized with PBS-BSA 3%-Triton 0.1% for 30 min. After a saturation step with PBS-3% BSA, the coverslides were incubated with Hoechst and Alexa Fluor™ Plus 647 Phalloidin for 60 min at room temperature. The cells were washed twice with PBS 1X and the coverslides were mounted onto slides and fixed with DAKO mounting medium. A Zeiss LSM880 confocal microscope was used to visualize up to 4 randomly selected fields per sample, taking z stacks of the whole sample thickness using. a Plan-Apochromat 40x/1.4 Oil DIC M27 lens. ImageJ software with the Fiji extension was used to process and analyse the images. For FACS analysis, cells were washed with PSB 1X, trypsinized, centrifuged and resuspended in PBS-FBS 1% at a final concentration of 5 x 10^5^ cells/mL The cell suspension was transferred to tubes and analyzed by flow cytometry (BD LSR-Fortessa). FlowJo software was used to process and analyze the data.

### Long-term exposure approach in vitro

MCF10A cells were chronically exposed to PS and PET (both 20 µg/mL) for 20 weeks, as single treatments or in combination with BMP_2_ (10 ng/mL). In parallel, passage-matched untreated cells and cells exposed to BMP2 and IL-6 (both 10 ng/mL) were also maintained. Cells were passaged twice a week, always replacing cell culture medium with new treatment-containing medium to ensure the constant exposure condition on the long-term. Functional assays and transcriptome and kinome analysis were performed at the end of the 20 weeks of exposure.

### Mammosphere formation assays

Cells were seeded at a density of 1500 cells/mL on 96-well ultra-low-attachment plates (Corning) in serum free basal mammary epithelial medium (Promocell) supplemented with B27, 20 ng/mL epithelial growth factor (EGF), 20 ng/mL basic fibroblast growth factor (bFGF), and 4 µg/mL heparin. Mammospheres were counted under a microscope using a 10X magnification after 6 days.

### Colony forming cell assays

Long-term exposed MCF10A were seeded under co-culture conditions with irradiated NIH-3T3 cells at a 1:128 ratio in the cell culture medium previously described. After 5 days, cells were fixed with methanol, stained with Wright Stain Solution, and washed repeatedly to remove excess of stain. Colonies were counted and classified based on their morphological features as luminal, myoepithelial or mixed colonies.

### Flow cytometry analysis of cell surface markers

Cells were seeded to reach 50-75% confluency on the day of the experiment. After trypsinization and PBS washes, the cells were double stained with: CD10 PE (ref 555375) / EPCAM FIT-C (ref 10109), CD44 PE (ref 550989) / CD24 FIT-C (ref 555427), CD24 PE (ref 559882) / CD49f (ref 555735); and the IgG control as appropriate: IgG1 MOUSE PE and FIT-C or IgG2a FITC. 2 µL of each antibody per 10^6^ cells was added to each well and incubated at 4ºC in the dark for 20 min. Then the cells were centrifuged at 1400 rpm for 5 min and supernatant was discarded. After a wash with PBS 1X, stained cells were resuspended in 300 µL of PSB-FBS 1% for analysis in a BD LSR-Fortessa analyzer. FlowJo software was used to process and analyze the data.

### Acini formation assays

3,000 cells/well were grown over a Matrigel coating (Corning) in µ-Slide 8 Well Glass Bottom (Ibidi). Cells were seeded in DMEM/F12 phenol red free media supplemented with 1% penicillin/streptomycin, 5% horse serum, 10 µg/mL insulin, 0.5 µg/mL hydrocortisone, 100 ng/mL cholera toxin, and 20 ng/mL EGF and 4% Matrigel. The medium was changed after 5 days and the cells were grown up to day 10. Then, the acini were washed twice with cold PBS (1x), fixed with PFA (4%) for 20 min, permeabilized with 0.2%Triton X100 for 30 min, washed three times with PBS-Glycine (0.1%), and saturated with PBS-BSA (3%). Nuclei were stained with Sytox Green Deep Red and actin was stained with PHA-rhodamine (both Thermo Fischer Scientific). After three washes with PBS, acini were ready for confocal imaging. Acini structural changes was visually scored and Image J was used to measure the area.

### Soft agar assay

Cell culture plate was coated with a bottom layer of 1.5% low-melting point agar (Promega) diluted in an equal volume of 2X culture medium to a final concentration of 0.75% and incubated at room temperature for 30 min. A suspension of 120,000 cells was added to 2 mL of 2X culture medium mixed in a 1:1 ratio with 0.9% low-melting point agar (0.45% as final concentration). Triplicates of 30,000 cells each were prepared by dispensing 1 mL of the mixture over the previously prepared agar layer. Once the agar was polymerized, the plates were incubated at 5% CO_2_ and 37ºC for 1 month before visual quantification of colony number each week.

### Migration and invasion assays

Cells were seeded at a density of 50,000 cells/well in Incucyte® Imagelock 96-well Microplate (Sartorius). The following day, the cells were treated with 5 µg/mL Mitomycin C (Merk) to stop proliferation. Wounds on the cell monolayer were created using the Incucyte WoundMaker, and after 2 washes with PBS, new medium was added. Continuous screening was carried out using the Live-Cell Analysis System Incucyte S3. Four to six technical replicates (wells) were seeded in each experiment. Wound size measurement was defined every 2 h for up to 24 h. The resulting wound healing data acquired by the Incucyte Scratch Wound Analysis Software Module was exported and analyzed. To evaluate cell invasiveness the plates were first coated with Matrigel at a 1/100 dilution in cell culture medium before adding a layer of 1/30 Matrigel dilution in medium on top of the cell monolayer after scratching. Wound size measurement was defined every 2 h for up to 72 h.

### Transcriptomic analysis

Three replicates of the long-term exposed MCF10A and the corresponding controls were seeded to reach approximately 80% confluency in the following 48 h. RNA from the sub-confluent culture was purified with RNeasy Plus Mini Kit (QIAGEN). The samples were sequenced by Novogene (UK) using the Illumina platform with a paired-end protocol and 150 nucleotides read length. Approximately 40 million reads per file were obtained and subsequently analyzed. R version 4.3.2 was used for statistical analyses through the R Studio software. Raw FASTQ files were quality filtered by Rfastp package, including sequence adapters removal, quality trimming and reads filtering when they were below 20 bases in length. Filtered reads were mapped to the human genome GRCh38 retrieved from GENCODE and counted using Rsubread package. Lowly expressed genes were automatically filtered by the filterByExpr function from edgeR package keeping as many genes as possible with worthwhile counts. calcNormFactors function from edgeR was used to calculate normalization factors between libraries. To guarantee that the observed differences were not due to technical artifacts, sva package was applied and inferred surrogate variants were included in the differential expression analysis performed by limma and voom packages. Differentially expressed genes (DEGs) were obtained by contrasting exposed samples to the negative control (CT).

To determine the enriched functional terms, over representation analysis (ORA) and gene set enrichment analysis (GSEA) were performed by clusterProfiler package. Two collections were used as models: Gene Ontology (GO) and MSigDB hallmark collection. Parameters were left by default except for maximum gene set size on GSEA which was set up to 800 and p values on ORA and GSEA, which were set up to 0.1 to get information of significant and close-to-significance values. Data visualization was performed by ggplot2 package. Raw and normalized data are available at the GEO depository under accession number GSE310440.

### KRT14 immunofluorescence

Cells (1500 per well) were seeded in 96-wells Phenoplates (Revvity) with the indicated treatments and grown for 48 hours and fixed with 4% formaldehyde for 15 minutes at room temperature (RT) before permeabilization for 3 minutes at RT with 0.2% Triton-X100 in PBS. After a 1 hour blocking step in PBS 5% BSA at RT, cells were incubated with a rabbit anti-human cytokeratin 14 (clone SP53 from ABCAM) primary antibody for 2 hours at RT in PBS 5%BSA, washed three times in PBS, incubated with an Dylight 488 goat anti-rabbit IgG secondary antibody (Invitrogen, 35552), Hoechst 33342 and Alexa Fluor 647 Phalloidin (Invitrogen, A30107) in the same buffer for 1 hour at RT and washed again 3 times in PBS. Labeled cells were imaged using an automated confocal microscope (Opera Phenix Plus, Revvity) at a 40x magnification. Images obtained after maximum projection of the z-stacks were analyzed using the Harmony software

### Kinome activity profiling

The effect of long-term exposure in kinases activity was evaluated. Three replicates of each experimental condition were seeded to reach 50 to 75% confluency on the day of collection. The cells were lysed on ice with M-PER (Mammalian Protein Extraction Reagent) lysis buffer mixed with 1/100 Protease Inhibitor Cocktail and 1/100 Phosphatase Inhibitor Cocktail (all Thermo Fischer Scientific). The lysates were centrifuged at 10,000 g and 4º for 10 min, and the collected supernatants were aliquoted, snap-frozen in liquid nitrogen and stored at -80ºC. Protein concentration was determined using the Bradford Protein Assay, with BSA (bovine serum albumin) as standard. The PamChip®12 Serine/Threonine (STK) and the PamChip®12 Tyrosine (PTK) peptide microarray systems (PamGene International B. V.) were used for kinome activity profiling, following the manufacturers’ protocol.

## Results

### Uptake of PS and PET Nanoplastics Modulates Human Stem Cell Self-Renewal and 3D Organization

The PS and PET NPLs used for this study differ in the type, size, and shape of polymer as indicated in Table S1. PET was obtained from the milling of bottles of water and corresponds to NPLs from a material frequently used in everyday life (*17*). In contrast, PS is of commercial origin and specifically produced for testing purposes. To determine whether PS and PET readily enter stem cells as reported for mature cells from various subtypes (*18-20*), we exposed CD34^+^ human hematopoietic stem cells isolated from healthy human bone marrow to fluorescent-labelled PS NPLs for the period of time indicated (Fig.1A). We observed a very rapid intra-cellular uptake of fluorescent-PS by CD34^+^ immature cells after 24 h of exposure. Comparatively, PS uptake by human primary healthy mammary cells, both immature (Epithelial SC) or mature (Fibroblasts), took slightly longer to accumulate within the cells (Fig.1B). Similar results were observed in a MCF10A-based model representing a series of early steps of breast cancer transformation (*7, 15, 21*) (Fig.1C). In all tested cell types, primary cells and cell lines, PS accumulated in the peri-nuclear region of the cell cytoplasm in lysosomal structures as identified by LAMP2 staining (Fig. S1A). We then compared the internalization of fluorescent-labelled PET and PS by the MCF10A epithelial SC cell line model (*7*). In both cases, we observed elevated peri-nuclear levels of PS and PET after 24 h (Fig. 1D). Interestingly, quantification of nanoplastics by flow cytometry revealed that cells internalized both particles within 3 h of treatment, and remained in the cytoplasm after 48 h (Fig. S1B). However, though PET internalization by cells had already reached its maximum at 3 h of exposure, with 100% of cells showing uptake, PS uptake remained below 60% even after 48 h, suggesting that PET is more readily internalized by cells than PS.

**Fig. 1.**
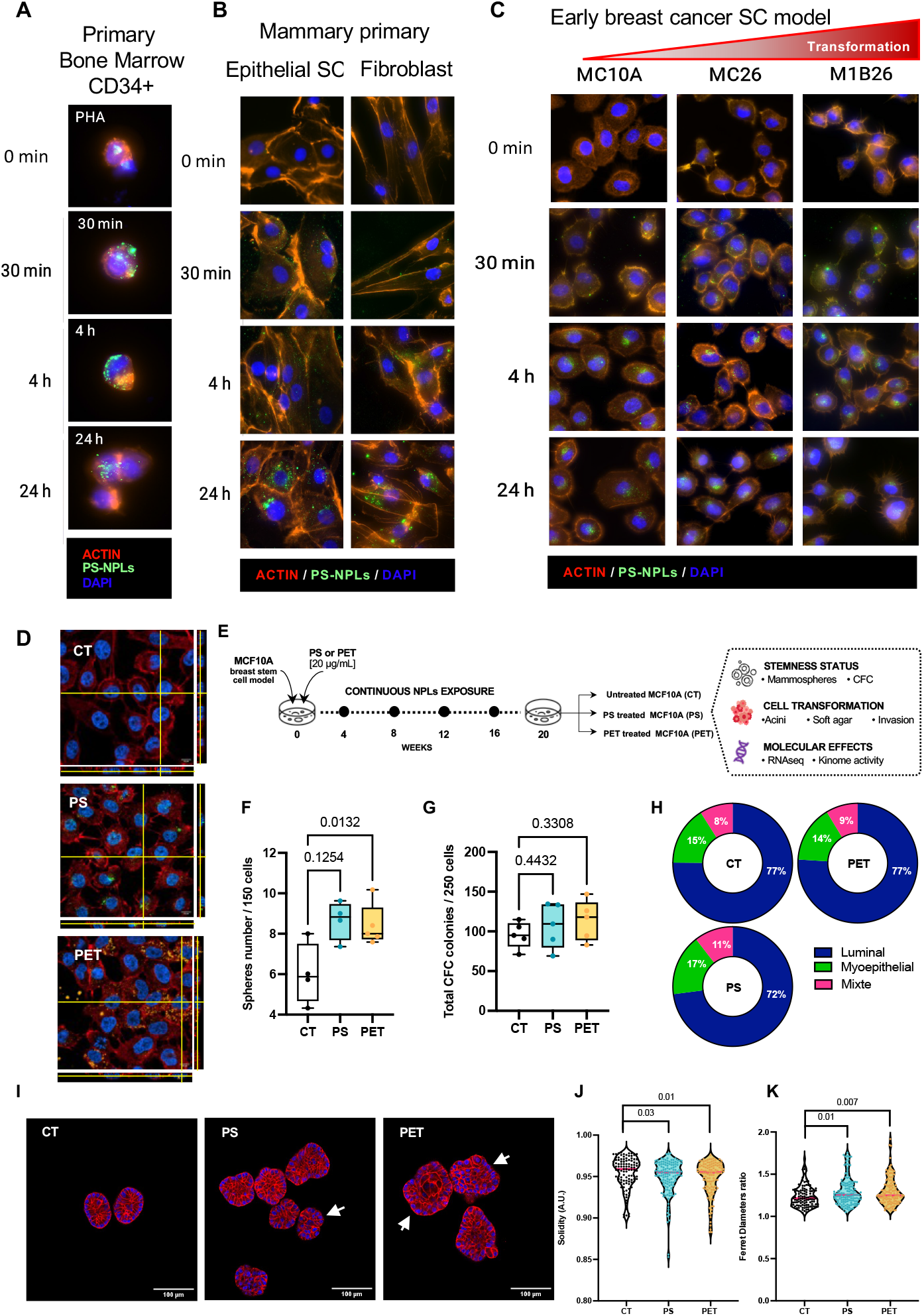
Long-term exposure of MCF10A cells to PS or PET NPLs affects their stemness and transformation status. Representative confocal images of fluorescent PS nanoplastic internalization after the indicated time in: (**A**) primary bone marrow CD34+ sorted hematopoietic immature cells or (**B**) epithelial or fibroblast cells isolated from primary mammary gland obtained from healthy donors or (**C**) MCF10A-derived models representing non-transformed mammary stem cells (MCF10A-CT), early (MC26) and more transformed (M1B26) cells as previously described (*7*). Nuclei are stained in blue, actin filaments in red, PS in green. (**D**) Confocal images of PS and PET nanoplastic internalization in MCF10A after 24 h compared to non-exposed controls. Nuclei are stained blue actin filaments red, PS green, PET orange. (**E**) Schematic diagram of the nanoplastic (NPLs) long-term exposure approach and analysis of their impact. (**F**) Quantification of the sphere-forming ability of untreated (CT) MCF10A cells or following 20 weeks of exposure to PS or PET nanoplastics (NPLs), n=5. (**G**) Quantification of total stem cell-derived progenitors by E-CFC assay (*15*), data represent the total number of colonies per 250 seeded cells, n=5. (**F**,**G**,) Each individual datum is represented. One way ANOVA is indicated on the graph by the P values. (**H**) Representation of the mean proportion of myoepithelial, mixed and luminal colonies quantified by E-CFC assay (n = 5). Data are expressed as the percentage of each subtype with a total number of scored colonies representing 100%. (**I**) Confocal images of acini. Nuclei are stained in blue, actin filaments in red, white arrows point to acini with a disorganized structure. Measurement of acini (**J**) solidity or (**K**) Ferret diameter ratio was quantified by image analysis using Fiji imaging software, n=109 to 131 individual acini. (**J**,**K**) Each individual data is represented. Mann-Whitney test is indicated on the graph by the P values.

To functionally explore the long-term impact of NPL uptake by stem cells, we continuously exposed MCF10A cells to PS or PET (20 µg/mL) for 20 weeks and then analyzed hallmark stem cell features, including self-renewal (sphere formation assay), multilineage differentiation (E-CFC progenitor assay) and 3D structure formation (acini formation assay) (Fig. 1E). Following 20 weeks of chronic exposure to either PS or PET, MCF10A sphere forming cell ability was significantly increased compared to untreated cells (Fig. 1F). However, their ability to differentiate was unaffected, as indicated by the absence of a significant effect on total E-CFC number (Fig. 1G), as well as their cell fate toward a luminal or myoepithelial lineage (Fig. 1H and Fig. S2A). Consistent with the SC expansion observed in the sphere-forming assay, NPL-treated MCF10A displayed altered acini structures (Fig. 1I). This was confirmed by a significant alteration of the morphological parameters that identified a decrease in the regularity of acini borders (Solidity, Fig. 1J). We also measured a significant increase in the Ferret diameters ratio (Fig.1K), corresponding to increased acini elongation (Fig. 1K), and a trend towards an increase in acini area (Fig. S2B).

Hence, PS and PET are rapidly taken up by immature human cells, stored in peri-nuclear structures and functionally impact stem cells features.

### PS and PET Nanoplastics Induce Mammary Stem Cells Transformation but Affect Distinct Cellular Pathways

Exposure to NPLs promotes the formation of larger, disorganized acini by human stem cells, suggesting morphogenetic alterations potentially linked to early transformation through distinct, material-specific mechanisms (*22*). To evaluate this hypothesis, we conducted a comparative study to assess the long-term effects of PS and PET NPLs on a human mammary epithelial stem cell MCF10A model, focusing on hallmark parameters associated with cellular transformation. Indeed, we quantified two of the most representative outcomes of cell transformation *in vitro*: the ability of cells to grow independently of anchorage (soft agar assay) and their invasive potential (Incucyte Matrigel invasion assay, previously associated with *in vivo* output (*7, 15*)). We observed that cells exposed to PS, and more significantly to PET, formed a greater number of colonies in the soft agar assay compared to non-exposed cells (Fig. 2A). We then assessed the invasive potential of cells with an adapted protocol based on the wound healing assay (Fig. 2B). Although the difference in wound healing was not statistically significant between control and NPL-treated cells, we observed a clear trend towards increased invading ability of NPL-exposed MCF10A (Fig. 2C). Remarkably, control cells required 65 h to close half of the wound while cells exposed to PS or PET took 38 and 41 h, respectively.

**Fig. 2.**
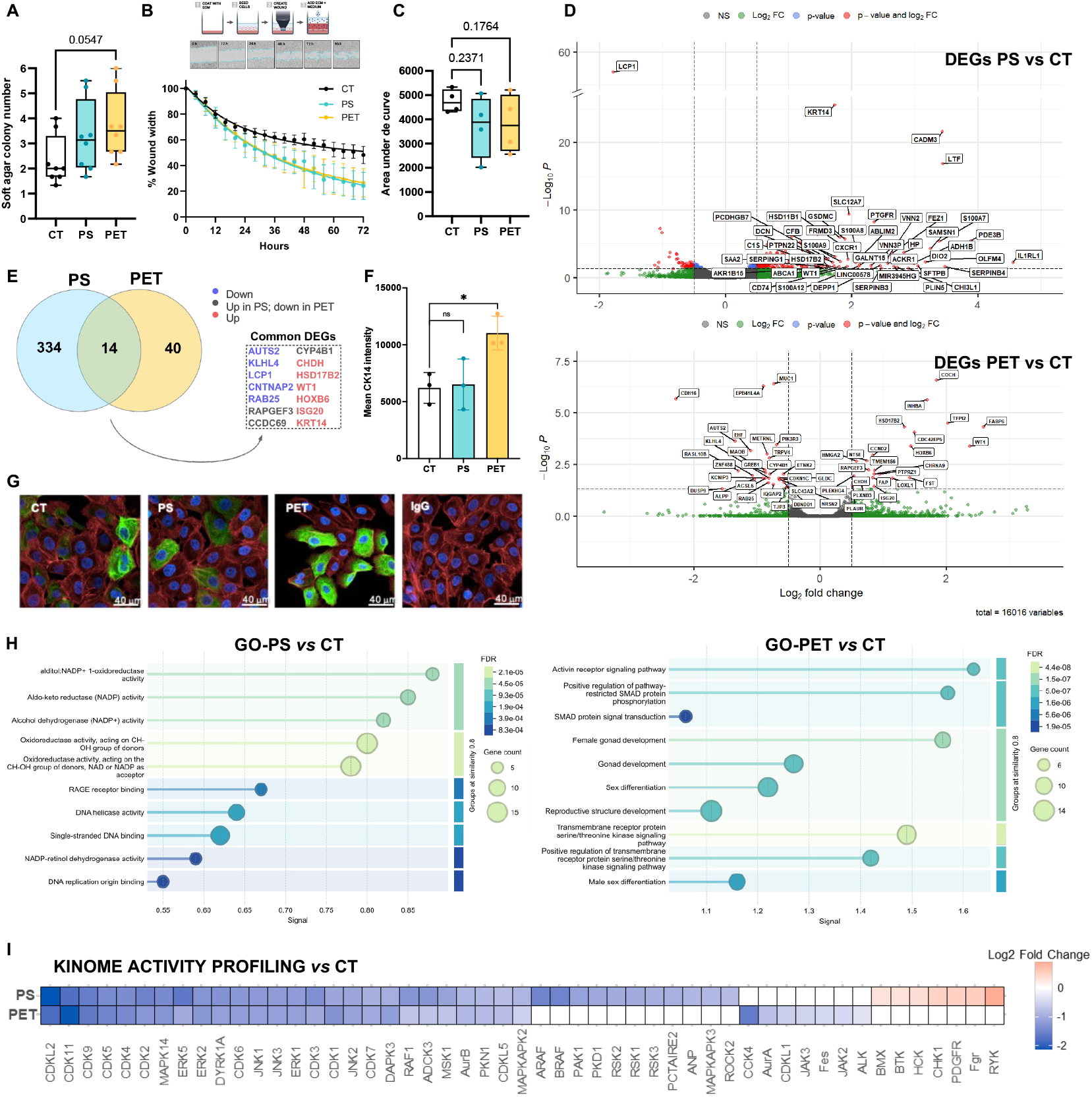
PS and PET nanoplastics differentially impact the transcriptome and kinome driving a preneoplastic stage. (**A**) Number of soft agar colonies formed after 20 weeks of exposure of MCF10A cells, n=8. (**B**) Schematic representation of the invasion assay and curve showing the kinetics of the wound width reduction and (**C**) quantification of the area under the curve measured after 72 h of treatment. Error bars represent the SEM, each point represents an independent *n=4*. (**D**) Volcano plot showing DEGs induced by PS (upper panel) and PET (lower panel) compared to negative controls. (**E**) Venn diagram of DEGs and list of the 14 common genes with those downregulated in blue, upregulated in red, and variable in grey. (**F**) KRT14 protein expression levels quantified (n=3) from immunofluorescent labelling of KRT14 as illustrated in (**G**) where KRT14 is labelled in green, actin in red and nuclei in blue. (**H**) GSEA analysis of the top 10 enriched terms from GO and Hallmarks databases. (**I**) Heatmap representation of differentially activated kinases in PS and PET exposed cells compared to non-exposed controls. Kinases with reduced activity are shown in blue, those with higher activity in red, and non-significantly changed kinases in white. (**A, C, F**) Significance measured using one-way ANOVA is indicated on the graph by the p-values.

Functional analyses revealed comparable qualitative effects between cells exposed to PS or PET, albeit the magnitude of the response was consistently greater with PET. To further investigate these differences, we performed transcriptomic profiling (RNA-seq) and kinase activation assays after 20 weeks of exposure. Differentially expressed genes (DEGs) commonly modulated by PS and PET were either upregulated (>0) or downregulated (<0) (Fig. 2D). A comparison across experimental conditions showed that PS and PET induced 348 and 54 DEGs, respectively, and identified a set of 14 commonly modulated genes (Fig.2E). Among the shared upregulated genes, *HOXB6* (Fig. S3A) and *KRT14* (Fig. S3B) expression was also higher during stem cell transformation in the MCF10A-derived breast cancer progression model (*7*). The upregulation of *KRT14*, a well-established marker of basal-like breast cancers, was further confirmed at the protein level by immunofluorescence staining in MCF10A cells chronically exposed to PS or PET (Fig. 2F, 2G). Importantly, the fact that only fourteen of these DEGs are shared between the two nanoplastics suggests distinct mechanisms of action for each type of NPL. This was confirmed by Gene Ontology analysis which revealed that (i) exposure to PS enhanced the cellular metabolism, detoxification processes and oxidative stress response pathways, but repressed biological functions aimed at preserving genome maintenance, such as DNA replication and DNA repair (Fig. 2F left panel), suggesting a potential impairment of cellular homeostasis, and (ii) long-term exposure to PET activated various receptor-associated pathways, including serine/threonine transmembrane kinases and SMAD-related signaling elements, which are typically triggered by receptors possessing serine/threonine kinase activity (Fig. 2F right panel). Using the MsigDB database for GSEA analysis further highlighted enriched pathways related to inflammatory and immune process, epithelial-to-mesenchymal transition and cell division following exposure to PS (Fig. S4 top), while PET significantly deregulated extracellular matrix organization pathways (Fig. S4 bottom).

Differences in protein kinase activity were also observed in exposed cells, as evidenced through a kinome assay. The kinase activity of 26 kinases were equally repressed by PS and PET, including those involved in cell cycle regulation (*e*.*g*., CDK1, CDK2, AurB) and stress-related signaling pathways (*e*.*g*., JNK, ERK, MSK1). However, the RSK family of kinases was specifically repressed by PS, while PET affected the activity of JAK kinases. Moreover, exposure to PS specifically activated a subset of kinases involved in critical cellular processes, including signal transduction (*e*.*g*., BMX, BTK, HCK, FGR, PDGFR, RYK), genome surveillance (CHK1), and proto-oncogenic signaling. Several of these kinases are known to be implicated in pathological conditions such as chronic inflammation, tumor progression, and metastasis, suggesting that PS may trigger molecular programs associated with disease development and malignant transformation (Fig. 2G).

Taken together, functional, transcriptomic and kinome analyses reveal that long-term exposure to NPLs could lead to early-stage transformation of mammary epithelial SCs by modulating distinct target genes or kinases, indicating that PS and PET may differ in their long-term effects.

### NPLs Cooperate with BMP2 to Drive Early Mammary Transformation

To further dissect the molecular consequences of long-term exposure to nanoplastics and determine whether PS and PET differentially engage molecular programs associated with early tumorigenesis, we extended our transcriptomic analysis to evaluate key signaling pathways. We applied single-sample Gene Set Enrichment Analysis (ssGSEA) using the Gene Ontology Biological Process (GOBP) and Hallmark collections to explore specific transcriptional signatures, focusing on inflammation, genomic instability, and DNA repair pathways.

Consistent with our other analyses (Fig. S4), we observed a trend toward enrichment of the inflammatory score following long-term exposure to PS, approaching statistical significance (p = 0.0660, Fig. 3A), similar to the pattern observed in the MCF10A-derived early breast cancer models (Fig. S5A). Among the genes included in the GOBP inflammatory response signature, we detected IL6, a cytokine that we previously identified as a key microenvironmental factor contributing to early stages of mammary epithelial cell transformation (*15*). While minimal differences were observed at the transcript level for a set of cytokines representative of inflammatory signals (Fig. S5B), MCF10A cells displayed a significant increase in IL-6 secretion, as measured by ELISA, upon 20 weeks of exposure to PS or PET (Fig. 3B). This confirms the impact of long-term exposure to these NPLs on the activation of inflammatory signals by epithelial SCs. Conversely, when we evaluated the impact on other major biological pathways, we observed no significant changes on genome instability (CIN70, Fig. S5C), DNA repair (Fig. S5D) or EMT (Fig. S5E) signaling using ssGSEA analyses.

**Fig. 3.**
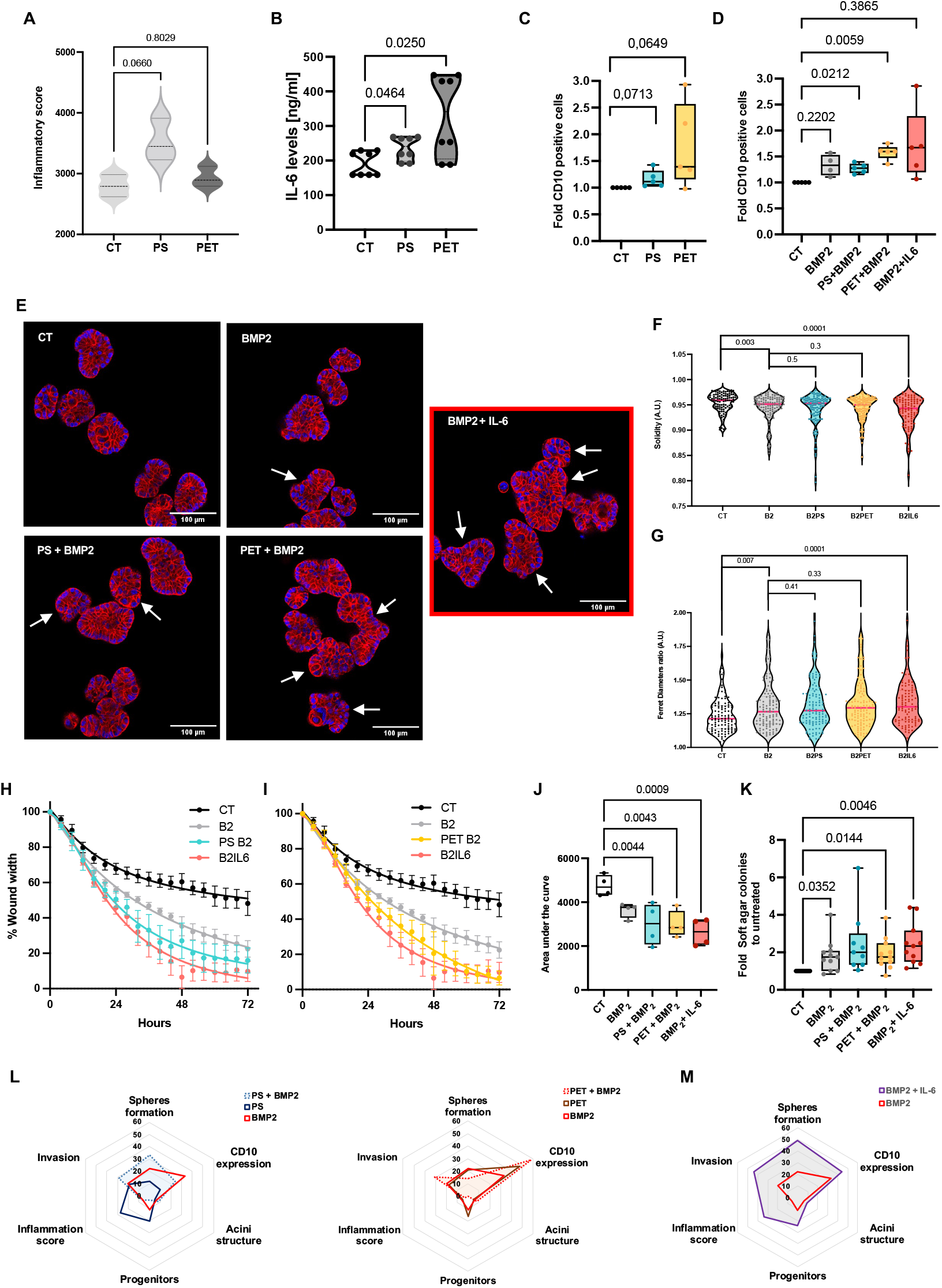
Co-exposure of epithelial stem cells to NPLs and BMP2 promotes transformation through different NPL-dependent mechanisms. (**A**) ssGSEA analysis of the inflammatory score in indicated RNAseq data, *n=3*. (**B**) Quantification by ELISA of IL6 in a 48-h culture supernatant of MCF10A cells following 20 weeks of exposure to PS or PET NPLs, n=8. (**C, D**) Flow cytometry quantification of CD10-cell membrane expression in cells treated by (**C**) PS or PET nanoplastics alone or (**D**) in combination with BMP2. Data are expressed as the fold increase in CD10-positive cells to untreated MCF10A cells. **(E**) Confocal images of acini nuclei stained in blue, actin filaments in red, white arrows point to acini with a disorganized structure. Measurement of acini (**F**) solidity or (**G**) Ferret diameter was quantified by image analysis using Fiji imaging software, n=109 to 131 individual acini. (**H**,**I**) Invasion assay curves showing the kinetics of the wound width reduction following 20 weeks of untreated (CT) or BMP2-treated MCF10A in the presence or not of PS (**H**) or PET (**I**). BMP2 and IL6 treatment is used as a positive control of the induction of a transformed state. (**J**) Quantification of the area under the migration curve, *n=4*. (**K**) Quantification of soft agar colonies. (**F, G, J, K**) Each individual datum is represented. One way ANOVA is indicated on the graph by the P values. (**L**) Radar plot representation of MCF10A cells features exposed for 20 weeks to PS (left panel), or PET (right panel) alone or in combination with BMP2. **(M)** Radar plot representation of MCF10A cells features exposed for 20 weeks to BMP2 or BMP2+IL6. Data represent the % of impact compared to untreated MCF10A cells. The functional and molecular readouts are indicated on the graph.

As shown with Venn diagrams, we uncovered genes jointly modulated by long-term plastic exposure (PS and PET) and genes previously identified as preneoplastic biomarkers in stem-like cells (ENI10 score) in our breast cancer progression model (*7*). Interestingly, while 28 genes were common to PS-treated MCF10A cells and the ENI10-signature, only one gene (FAP) was expressed by PET-exposed cells (Fig. S5F). ssGSEA enrichment score analysis using the 29-gene list (designated ENI10-NP) revealed modulation of this score following treatment with the plastics, displaying a pattern similar to that observed in the MCF10A-derived breast cancer model (Fig. S5F). Moreover, we observed an increase in cells expressing the CD10/MME cell surface marker (enriched in normal or stem-like cells (*23*)) (Fig. 3C), as well as stem cell expansion (sphere-forming ability) (Fig. 1F) following treatment with both NPLs. This supports the impact of NPLs long-term exposure on stem cell expansion as sustained by increase in both spheres forming ability and CD10-membrane positive cells.

Our previous findings on the role of BMP2 and IL-6 in the early transformation of epithelial SCs unveiled the activation of SMAD signaling components (*7, 15*), which were also activated herein upon exposure to PET, as evidenced in our transcriptomic analysis (Fig. 2F, right panel). This suggests that the BMP2 pathway may already be primed in these cells as reported for another environmental contaminant Bisphenol A (*16*). We then tested whether chronic co-exposure to NPLs and BMP2 could synergize to promote neoplastic progression and assessed the carcinogenic potential of combined long-term exposure to NPLs and BMP2 including BMP2+IL6 co-treatment as a positive control of transformation (*7, 15*). Importantly, we obtained a similar increase in the levels of CD10-positive cells upon treatment with PET for 20 weeks alone, as with the transformation-positive control BMP2+IL6 (Fig. 3D). Of note, we found no significant difference by co-exposing cells with BMP2 in terms of cell division doubling time (Fig. S6A), sphere-forming ability (Fig. S6B) and total progenitor differentiation (Fig. S6C). Acini formation in the presence of BMP2 was more strongly affected in both size and shape (Fig. 3E) compared to NPL treatment alone (Fig. 1I), as illustrated by solidity (Fig. 3F) and Ferret diameter (Fig. 3G) measurements. Notably, the combination of BMP2 and NPLs induced effects comparable to those observed in the BMP2+IL-6 transforming condition. Co-exposed cells formed larger and more structurally disorganized acini, particularly in the PET+BMP2 group, which closely mirrored the disorganization seen with BMP2+IL-6.

Finally, we measured the cell motility (Fig. S6F), invasive potential and ability to form clones in soft-agar of cells treated with PS+BMP2 or PET+BMP2 to directly evaluate the transformation state. We observed a statistically significant impact of BMP2 cooperation with both NPLs that drove cell invasion (Fig. 3H-J). Moreover, our soft agar results provided the most compelling evidence of transformation-like behavior as NPL- and BMP2-treated cells exhibited a marked increase in anchorage-independent growth (Fig. 3K). Both assays indicate values for NPLs+BMP2 comparable to the transforming BMP2+IL-6 condition. Ultimately, both NPLs led to biological effects consistent with cellular transformation as supported by different functional evaluations compiled and illustrated on a radar plot representation to compare the global effects of PS, PET and BMP2 combined or not on epithelial stem cells (Fig. 3L), compared to the established transformed BMP2+IL6 condition (Fig. 3M). These data integration highlight that, inversely to PS and BMP2, PET and BMP2 have very similar functional impacts.

Altogether, these findings underscore the synergistic effect of NPLs and BMP2 in promoting malignant traits and support the notion that exposure to NPLs drives epithelial SCs toward a neoplastic state, especially under microenvironmental conditions that mimic early tumorigenesis.

### NPLs and BMP2 Cooperate to Promote a Preneoplastic State in Epithelial Stem Cells through distinct Mechanisms while Sharing a Unique Biomarker Risk Signature

Next, to identify molecular effectors implicated in the functional response of MCF10A cells exposed to NPLs and BMP2 or not, we analyzed their transcriptomic profiles after a 20-week induction. Cells co-exposed to PET+BMP2 harbored more dysregulated genes (778 DEGs) than PS+BMP2 (368 DEGs), compared to untreated MCF10A cells (Fig. 4A). We identified 78 genes common to the transforming positive control condition (BMP2+IL-6) and PS+BMP2, and 294 genes for PET+BMP2 treatments. Co-exposure to PET and BMP2 modulated the expression of 461 genes compared to BMP2 alone, whereas PS combined with BMP2 affected only 138 genes. Despite this difference in gene modulation, over representation analysis (ORA) of DEGs did not reveal any transcriptionally prominent pathway uniquely impacted by the BMP2+PS combination (Fig. 4B, upper panel), suggesting that PS does not significantly influence the transcriptional effects induced by BMP2 at this level of analysis. However, it is worth noting that exposure to PS enhanced the impact of BMP2 on actin filament organization and pathways related to ROS production. These processes are involved in breast tumor initiation and progression, particularly through cell migration, plasticity or oxidative stress and DNA damage. In contrast to PS, PET combined with BMP2 elicited a markedly broader transcriptional response, significantly increasing the expression of genes involved in stress-activated signaling, RHO GTPase, NF-κB, and RAS protein signal transduction, none of which were affected by BMP2 alone (Fig. 4B, lower panel). These pathways are known to contribute to breast cancer progression through their roles in cell migration, inflammation, and survival. Interestingly, however, when we assessed the long-term effects of PS and PET in cooperation with BMP2 on kinome activity, PS+BMP2 emerged as the condition most closely mimicking the BMP2+IL-6 positive control. Notably, we observed the upregulation of 8 kinases also upregulated in the BMP2+IL-6 condition. The upregulation of BMX, Src, Lyn, Abl, Fyn, HCK, RYK, and PDGFR in response to PS treatment highlights a kinase activation profile that is particularly relevant to the basal-like breast cancer subtype. In contrast, PET+BMP2 predominantly downregulated kinases activity, with a profile largely overlapping that of BMP2 alone (Fig. 4C). Altogether, these analyses confirm distinct modes of action for the two NPLs, with PET+BMP2 exerting a broader impact at the transcriptomic level, while PS+BMP2 primarily activates a specific subset of kinases. Despite these mechanistic differences, both NPLs ultimately led to biological effects consistent with cellular transformation, particularly in the context of the basal-like breast cancer subtype, as supported by functional evaluations (Fig. 3).

**Fig. 4.**
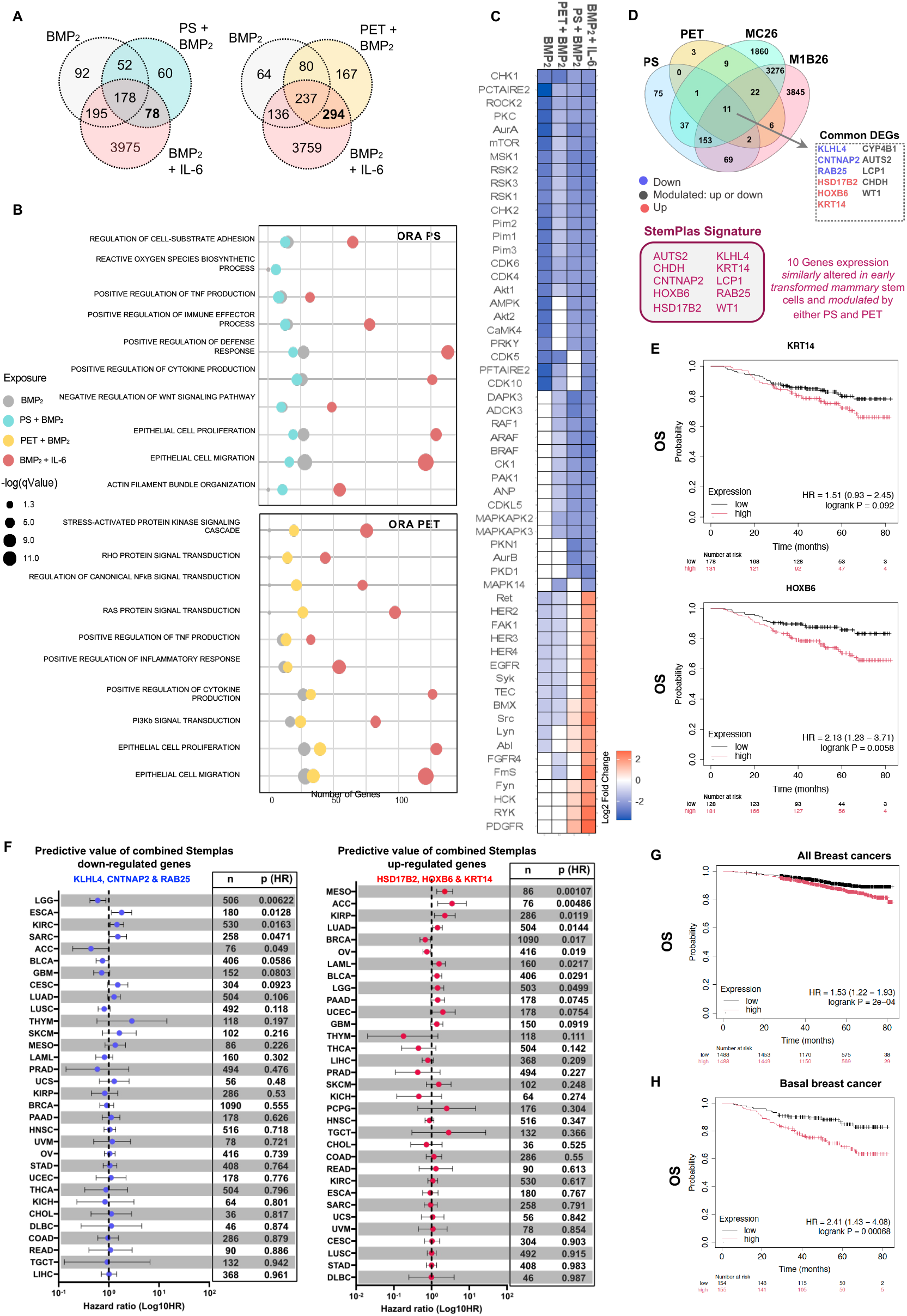
Transcriptome and kinome analysis reveal different mechanisms of SC transformation upon NPL and BMP2 exposure, associated with a molecular signature (StemPlas) predictive of breast cancer patient outcome. (**A**) Venn diagram representation comparing the DEGs for PS+BMP2 (left) and PET+BMP2 (right) with the DEGs identified in controls (BMP2 and BMP2+IL-6). (**B**) ORA analysis showing the 10 most relevant GO terms enriched after long-term co-exposure to PS+BMP2 (top) or PET+BMP2 (bottom). (**C**) Hierarchical heatmap representation of differentially-activated kinases in PS+BMP2, PET+BMP2, BMP2, and BMP2+IL-6 exposure conditions compared to non-exposed controls. Kinases with reduced activity are represented in blue, those with higher activity in red and non-significantly changed kinases are shown in white. (**D**) Venn diagram comparing the DEGs for PS, PET and established transformed MCF10A (MC26 and M1B26) (*7*). (**E**) Kaplan Meier analysis of overall survival (OS) of basal breast cancer patients stratified by KRT14 (up) or HoxB6 (bottom) expression levels. (**F**). Correlation of the genes identified as commonly down-(blue, left panel) or up-(red, right panel) regulated upon exposure to NLPs or transformation (see panel 4D) and survival outcome for the indicated types of cancers of the TCGA Pan-Cancer atlas estimated by hazard ratios of progression-free survival corresponding to one standard deviation of the score taken as a continuous variable. Dots show the hazard ratio and adjacent bars the 95% confidence interval. Tumor samples from the TCGA’s Pan-Cancer atlas linked by grey lines and the difference are color coded on the dots representing the tumor samples. ACC: adrenocortical carcinoma, BLCA: bladder urothelial carcinoma, BRCA: breast invasive carcinoma, CESC: Cervical Squamous Cell Carcinoma and Endocervical Adenocarcinoma, CHOL: cholangiocarcinoma, COAD: colon adenocarcinoma, DLBC: Lymphoid Neoplasm Diffuse Large B-cell Lymphoma, ESCA: esophageal carcinoma, HNSC: head and neck squamous cell carcinoma, KICH: kidney chromophobe, KIRC: kidney renal clear cell carcinoma, KIRP: kidney renal papillary cell carcinoma, LAML: acute myeloid leukemia, LGG: low-grade glioma, LIHC: liver hepatocellular carcinoma, LUAD: lung adenocarcinoma, LUSC: lung squamous cell carcinoma, MESO: mesothelioma, OV: ovarian cancer, PCPG: pheochromocytoma and paraganglioma, PRAD: prostate adenocarcinoma, READ: rectum adenocarcinoma, SARC: sarcoma, SKCM: skin cutaneous melanoma, STAD: stomach adenocarcinoma, TGCT: testicular germ cell tumor, THCA: thyroid carcinoma, THYM: thymoma, UCEC: uterine corpus endometrial carcinoma, UCS: uterine carcinosarcoma, UVM: uveal melanoma. (**G, H**) Kaplan Meier analysis of overall survival (OS) stratified according to high or low StemPlas expression based on median transcript levels in all types of breast cancer (**G**) or basal breast cancer (**H**).

To determine early biomarkers of cancer risk through stem cell transformation owing to plastic exposure, we compared RNAseq transcriptomic data obtained from MCF10A exposed over the long-term to NPLs to our MCF10A-derived breast cancer progression model (*7*). This allowed us to identify a specific molecular signature associated with the exposure of our mammary stem cell model to plastic particles, including genes commonly dysregulated at an early stage of MCF10A transformation (*7, 15*). We identified a set of 10 genes including transcripts up- or down-modulated by both NPLs (constituting the StemPlas signature), and specifically affected by plastic exposure (Fig. 4D). Several of these markers are part of transcriptional programs associated with breast cancer progression, with some more particularly dysregulated in the basal-like subtype, such as *KRT14*. Analysis of public RNAseq datasets using Kaplan-Meier survival curves stratified according to gene expression levels, confirmed that high *KRT14* expression is associated with significantly poorer overall survival (OS) in patients with basal breast cancer as expected (Fig. 4E, upper panel). Impressively, a highly significant association was also found for *HOXB6*, a transcription factor commonly upregulated by both types of NPLs and frequently overexpressed in cancer, including breast cancer (Fig. 4E, lower panel). These findings reinforce the hypothesis that plastic-induced reprogramming likely activates early oncogenic pathways in the SC compartment. We analyzed the ability of NPL-dysregulated genes, three downregulated (Fig. 4F left panel) and three upregulated (Fig. 4F right panel)) included in the StemPlas signature (Fig. 4D) to predict patients’ survival in 33 different cancers in the TCGA database (*24*). Analysis by Cox regression models highlighted that the transcripts levels of these limited gene set are however able to statistically predict poor overall survival (OS) patient outcome in various other cancer types. The most predictive values were obtained for low grade glioma (LGG), bladder urothelial carcinoma (BCLA) and adrenocortical carcinoma (ACC). Lastly, when evaluating the relevance of the defined StemPlas signature in breast cancer patients regardless of subtype, we observed significantly poorer overall survival in the group with high StemPlas expression levels (Fig. 4G). This molecular signature also strongly predicted outcome specifically in basal breast cancer patients (Fig. 4H). Indeed, high StemPlas expression was significantly associated with reduced overall survival, suggesting its potential role as a prognostic marker in breast cancer.

Together, these findings highlight the prognostic value of the StemPlas signature and suggest that chronic exposure to NPLs may establish a cellular context conducive to stem cell transformation, thereby contributing to aggressive tumor phenotypes, particularly within the basal breast cancer subtype.

## Discussion

The omnipresence of nanoplastics (NPLs) in the environment has raised growing concerns about their long-term impact on human health. While most studies to date have focused on short-term exposure to pristine polystyrene (PS) nanoplastics, our work addresses a critical gap by modeling chronic, low-dose exposure using environmentally-relevant NPLs, namely PS and polyethylene terephthalate (PET), in a stem cell (SC) context. We followed this approach to more accurately reflect real-world exposure scenarios, where individuals are continuously exposed to a mixture of plastic particles with diverse physicochemical properties as illustrated in our study by using PS and PET.

In this context, it is urgent to evaluate the impact of long-term exposure on adult residual tissue stem cells as previously suggested (*10, 14, 25*). Indeed, stem cells constitute a reservoir of cells able to accumulate alterations induced by chronic exposure, which may eventually initiate the carcinogenic process (*26*). The detection of NPLs in human breast milk (*11, 12*) strongly suggests that the mammary gland is a secondary target organ of exposure, particularly vulnerable due to its sensitivity to endocrine-disrupting chemicals present in plastic polymers and NPLs (*13, 15, 16*). In this work we have characterize the effects of long-term NPLs exposure on MCF10A, an epithelial stem cell line model, also known as a robust model to evaluate transformation ability (*7, 15, 16, 21, 32*).

We demonstrated here that chronic exposure to NPLs, particularly in the presence of BMP2, a key regulator of mammary stem cell fate and a known contributor to breast cancer initiation and progression (*15*), can reprogram mammary epithelial cells toward a pro-oncogenic state. BMP2 is an intrinsic component of the mammary microenvironment, and its dysregulation has been implicated in early tumorigenesis (*8, 16, 33*). Functionally, both PS and PET promoted the expansion of the epithelial stem cell compartment, as evidenced by increased mammosphere formation, and disrupted tissue architecture in 3D acini assays. While BMP2 co-exposure did not further enhance sphere-forming capacity, it markedly exacerbated acinar disorganization, an early hallmark of transformation, highlighting a cooperative effect between NPLs and endogenous BMP signaling. Although PS and PET alone induced trends toward increased anchorage-independent growth and migration, these effects did not reach statistical significance. However, in the presence of BMP2, these phenotypes were significantly amplified, supporting a functional synergy between plastic-derived particles and BMP2 in driving early oncogenic events.

At the molecular level, exposure to PS led to the upregulation of a kinase network including BMX, Src, Lyn, Fyn, Abl, HCK, RYK, and PDGFR, many of which are associated with basal-like breast cancer, a subtype characterized by poor prognosis and limited therapeutic options. These kinases are known to promote epithelial-to-mesenchymal transition (EMT), invasion, and metastasis, and their activation in cells co-exposed to PS and BMP2 mirrors the transforming profile observed with BMP2 in an inflammatory context (6), suggesting that PS may potentiate BMP2-driven oncogenic signaling through inflammatory and mesenchymal reprogramming. In contrast, PET primarily induced transcriptional dysregulation, highlighting distinct but converging mechanisms of action between different NPLs. Interestingly, PET and BMP2 appeared to act in a more cumulative rather than complementary manner, suggesting a different mode of cooperation compared to PS. Despite these mechanistic differences, both NPLs ultimately led to similar phenotypic outcomes, including the upregulation of Keratin 14 and the emergence of a specific molecular signature, which we referred to as StemPlas, comprising 6 genes. Importantly, a broad analysis of the TCGA database using this limited gene set revealed that it was predictive of poor overall survival in cancer patients, reaching statistical significance in patients with low-grade glioma (LGG), bladder urothelial carcinoma (BLCA), and adrenocortical carcinoma (ACC). Moreover, the entire StemPlas signature could predict poor overall survival in patients with breast cancer (BRCA), in particular for the basal subtype. From a mechanistic point of view, it is important to note that despite their distinct anatomical origins, basal-like breast cancer, LGG, BLCA, and ACC share several molecular and cellular features that contribute to tumor initiation and progression. These include frequent mutations in tumor suppressor genes, such as TP53 and CDKN2A/B, and dysregulation of key oncogenic signaling pathways including PI3K/AKT/mTOR, MAPK/ERK, and Wnt/β-catenin. All four tumor types exhibit oxidative stress and DNA damage responses, promoting genomic instability, as well as inflammatory signatures marked by NF-κB pathway activation and immune cell infiltration. Moreover, they display stem cell-like characteristics and cellular plasticity, which are associated with aggressive behavior and poor prognosis. Previously published transcriptomic and epigenetic analyses revealed overlapping gene expression programs related to proliferation, migration, survival, and immune modulation, suggesting convergent oncogenic mechanisms across these malignancies.

This supports the fact that the StemPlas signature may serve as a biomarker of plastic-induced transformation, and could be used to better understand the effects of environmental NPL pollutants on stem cell-related carcinogenesis. Indeed, while previous studies have shown that PS nanoplastics promote proliferation and adipogenic differentiation of mesenchymal stem cells (*27*), and that PET microplastics induce senescence and impair differentiation (*28*), the pro-carcinogenic effects of NPLs on stem cells have so far remained unexplored. Some studies suggested the role of NPLs in promoting breast carcinogenesis, as illustrated by the fact that polypropylene microplastics promoted the metastatic potential of MDA-MB-231 and MCF-7 cells (*29*). Nano- and micro-sized PS were shown to induced mild stimulatory effects of proliferation and migration of MDA-MB-231 and M13SV1 cells (*30*). Other types of plastics have also been reported as EMT-inducing in breast cells (*29*) and other target cell lines (*31*). Moreover, other environmental pollutants have also been implicated in carcinogenesis, such as (i) pesticides and air pollutants, linked to a higher risk of developing glioma, (ii) aromatic amines and polycyclic hydrocarbons implicated in bladder carcinogenesis, and (iii) endocrine-disrupting chemicals, such as bisphenol A and phthalates, that interfere with adrenal steroidogenesis in ACC. These findings reinforce the relevance of our model, which integrates chronic exposure to plastic-derived particles with stem cell transformation and microenvironmental signaling, and suggest that the StemPlas signature may reflect a broader environmental carcinogenesis mechanism.

Our findings underscore the urgent need to refine current approaches for evaluating the health risks associated with nanoplastics (NPLs). While PS and PET differ in their molecular mechanisms, PS primarily activating kinase networks and PET inducing transcriptional deregulation, both lead to similar oncogenic outcomes, particularly when combined with microenvironmental cues such as BMP2. This convergence highlights the importance of considering not only the chemical diversity of NPLs but also the cellular and tissue context in which exposure occurs. Importantly, our study reveals that BMP2, a key regulator of stem cell fate and a known target of other environmental pollutants like bisphenols, significantly amplifies the transforming potential of NPLs. This functional synergy suggests that chronic, low-dose co-exposure to multiple environmental agents may pose a greater carcinogenic risk than previously acknowledged. By demonstrating how NPLs interact with endogenous signaling pathways to promote stem cell transformation, our results provide a critical foundation for the development of more predictive and biologically-relevant testing platforms. Current risk assessment models often rely on short-term exposure to uniform, commercially available nanoplastics, which fail to capture the complexity of real-world scenarios. Our data advocate for the integration of stem cell-based assays, microenvironmental factors, and realistic NPL formulations into standardized testing frameworks. Such improvements are essential to accurately assess long-term health risks and to inform public health policies aimed at mitigating the impact of environmental nanoplastics.

## Acknowledgments

From CRCL platforms, we thank P. Battiston-Montagne and C. Vanbelle (PIC cytometry, imaging).

## Funding

This work was funded by:

European Union’s Horizon 2020 research and innovation programm PlasticHeal project under grant agreement No 965196 (VMS, AH); funding of post-doctoral fellow IB.

French Fundation for Research Medical-FRM, (team FRM EQU202203014695)(BG,VMS) Patients Associations: “Déchaîne Ton Cœur”; “Comité féminin pour le dépistage du cancer du sein 74” (VMS).

RUBAN ROSE Award Avenir 2021 (VMS), funding of doctoral fellow LB.

## Author contributions

Conceptualization: IB, BG, VMS

Methodology: IB, KG, LB JG, RE, LR, AH, BG, VMS

Investigation: IB, KG, BG, VMS

Visualization: IB, KG, JG, RE, BG, VMS

Funding acquisition: AH, BG, VMS

Project administration: BG, VMS

Supervision: BG, VMS

Writing – original draft: IB, VMS

Writing – review & editing: IB, BG, VMS

## Competing interests

Authors declare that they have no competing interests.

## Data and materials availability

Sequencing data have been deposited at GEO (GSE310440).

## GLP Compliance

All in vitro experiments were conducted in accordance with OECD GLP principles, using standardized protocols for nanoplastic preparation, exposure, and data handling.

## Notes

### Competing Interest Statement

The authors have declared no competing interest.

