## Supplemental Figures and tables for "Chronic Exposure to Nanoplastics Alters Stem Cell Type-Specific Mechanisms, Promoting Cancer Development"

**A**

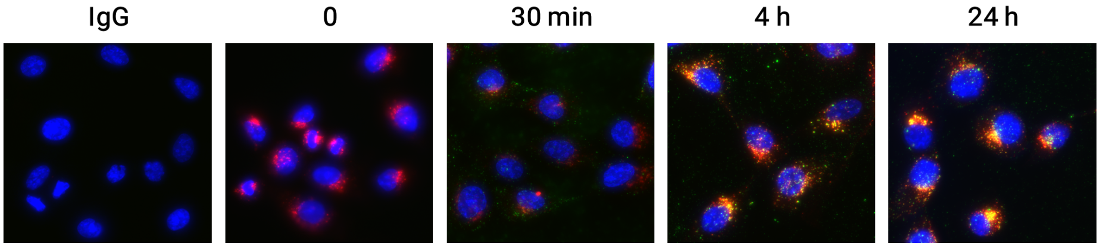


**B**

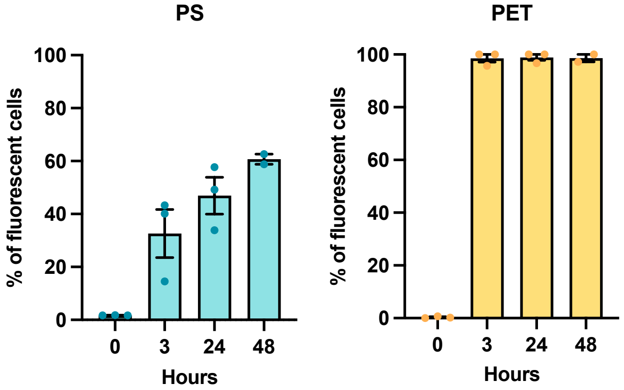


**Fig. S1. A.** Colocalization with the LAMP2 cell trafficking and lysosome marker of Fluorescent PS-GFP following the exposure indicated time and as compared to negative antibody control (IgG) Illustrative confocal images. **B.** Quantification of PS and PET nanoplastic (100 µg/mL) internalization in MCF10A cells represented as frequence of fluorescent cell in flow cytometry analysis.

A

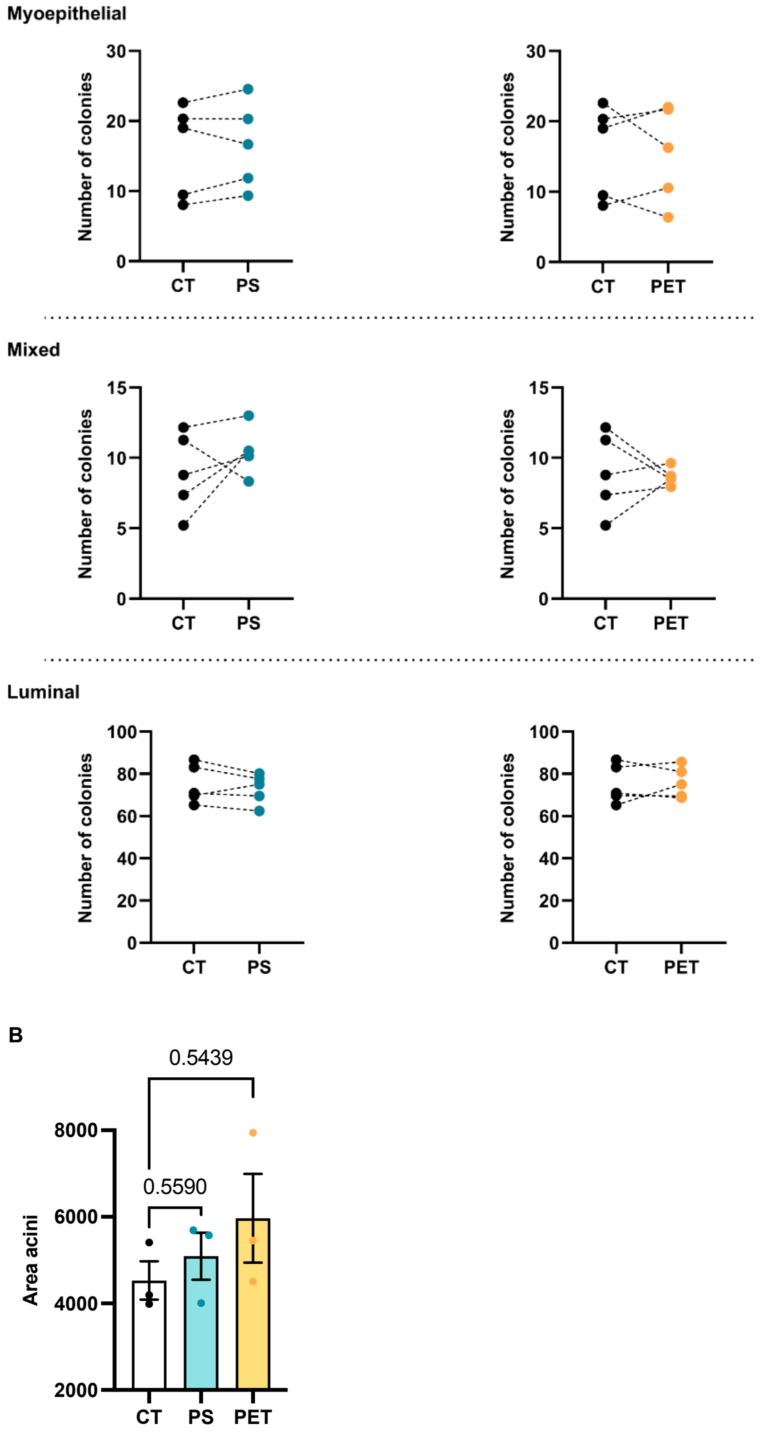


**Fig. S2. A.** Epithelial colonies forming cell quantification. Paired representation of the quantification of myoepithelial, mixed and luminal colonies in CFC assay (n = 5) comparing each type of colonies in non-exposed cells (CT) and after PS or PET long-term exposure. **B.** Acini Area measurement. One-way Anova test is indicated on the graph by the P values.

A

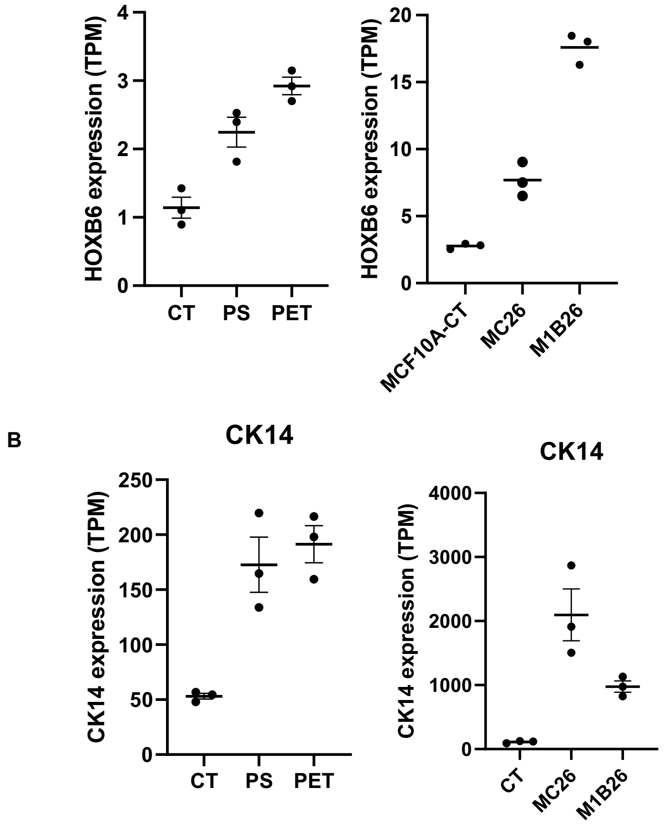


**Fig. S3.** Quantification of HOXB6 (**A**) and KRT14 (**B**) genes expression in long-term exposed MCF10A to PS or PET NPLs as compared to non-exposed controls (CT) or in the MCF10A early breast cancer transformation models (MC26 and M1B26) and their relative control (MCF10A-CT).

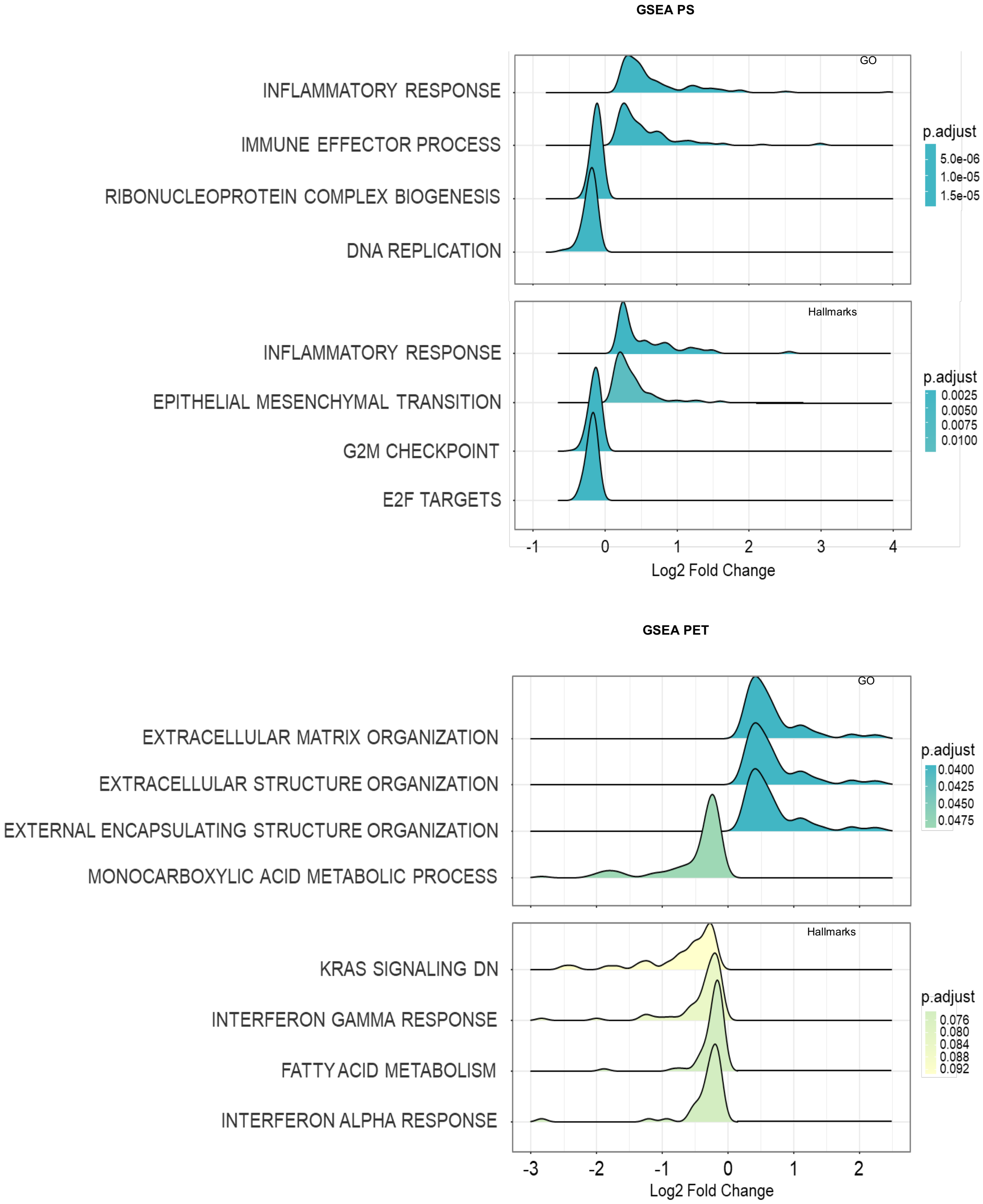


**Fig. S4.** GSEA analysis of the top 4 enriched terms from GO and Hallmarks databases in MCF10A cells exposed to PS (upper panel) or PET (lower panel) NPLs when compared to untreated control.

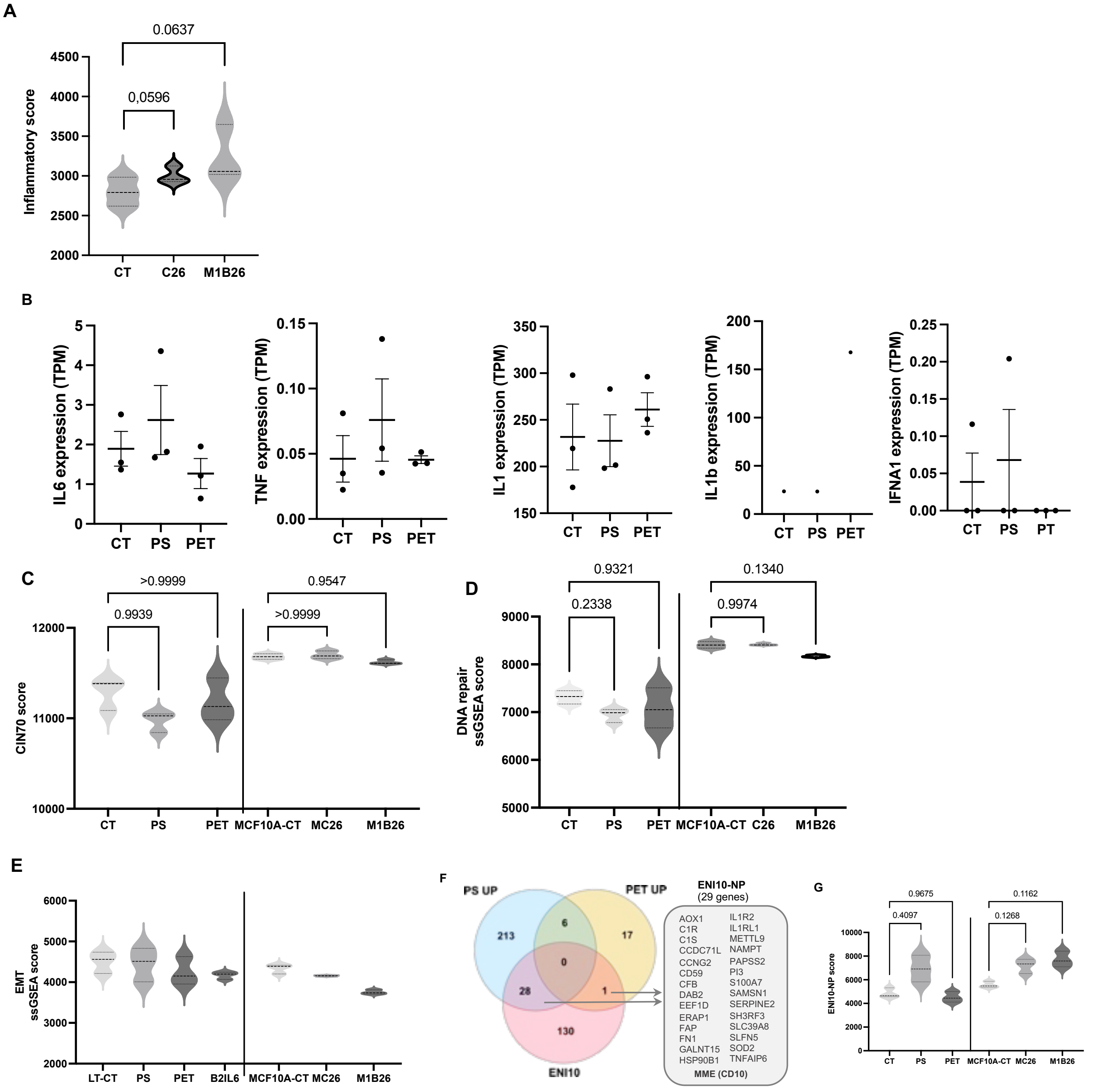


**Fig. S5. A.** ssGSEA analysis of inflammatory score of in indictaed RNAseq data, *n=3*. **B.** Expression of the indicated inflammatory cytokines in RNAseq data from cells untreated (CT) or exposed to PS or PET NPLs. **C.** ssGSEA analysis of CIN70 score in MCF10A exposed to PS or PET NPLs as compared to non-exposed controls (CT) or in the MCF10A early breast cancer transformation models (MC26 and M1B26) and their relative control (MCF10A-CT) *n=3*. **D.** As in C. for the DNA repair ssGSEA score. **E.** As in C. for the EMT ssGSEA score. **F**. Venn diagram representation of the intersection between genes modulated by either PS or PET NPLs and genes list (ENI10) identified as a hallmark of pre-neoplastic stage of cancer stem cells in a broad range of tumors. **G.** As in C. for the ENI10-NP ssGSEA score, a restrained 29-genes ENI10-NP derived signature.

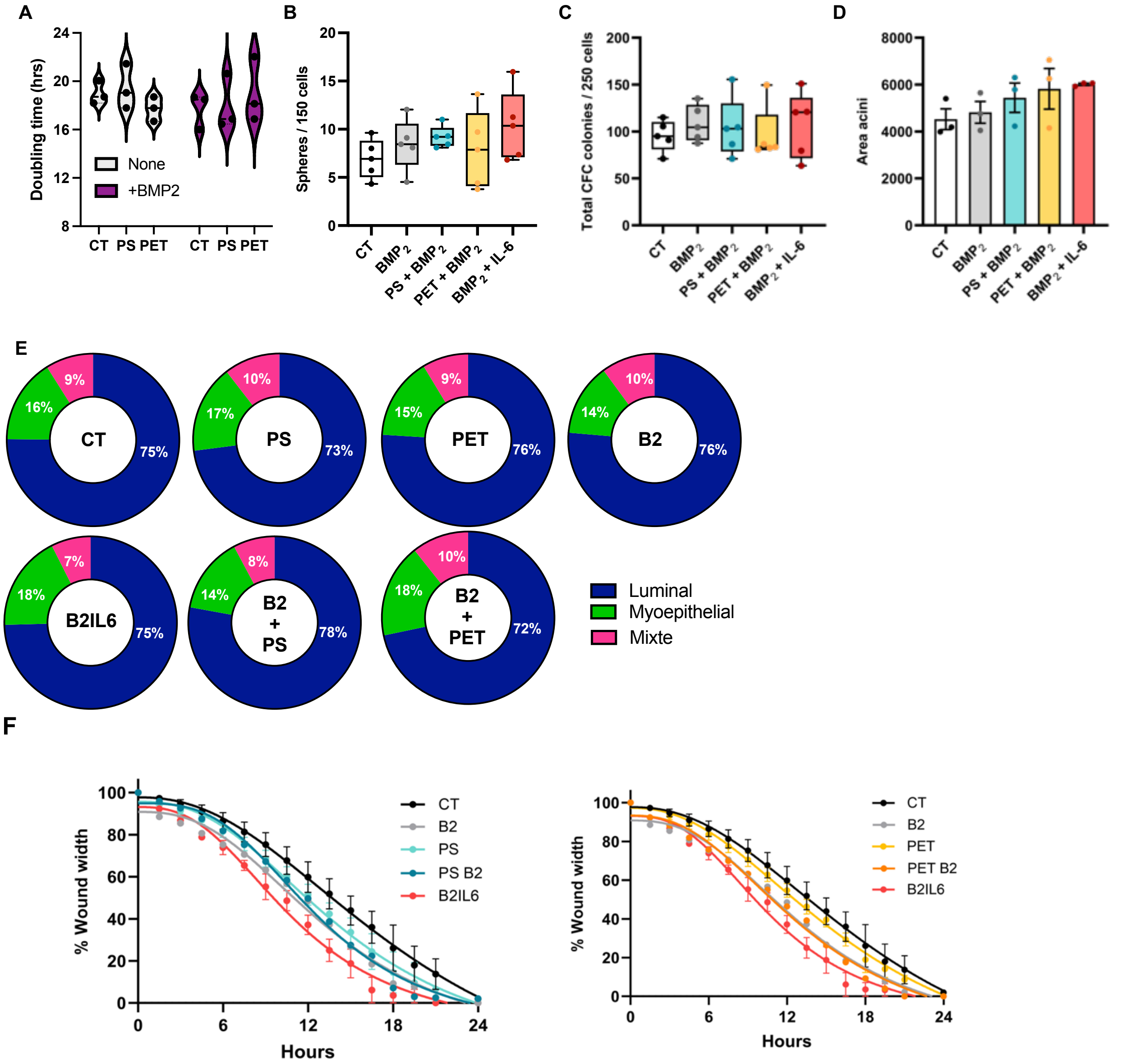


**Fig. S6. A**. Doubling time of each cellular models chronically cultivated 20 weeks in the presence of the indicted plastic with addition or not of BMP2. n=3. **B**. Spheres forming ability quantification comparing the effect of the different exposure conditions (BMP2 alone or in presence of PS, PET or IL6) to the non-exposed controls (n=5). **C**. Quantification the total colony number in E-CFC assay (n=5) (BMP2 alone or in presence of PS, PET or IL6). **D**. Average acini area in the indicated conditions (n=3). **E**. Quantification of myoepithelial, mixed and luminal colonies in E-CFC assay (n = 5) in the different exposure conditions indicated. Data are expressed as percentage of repartition for each colonies subtype with a total number of scored colonies representing 100%. **F**. Migration assay curves showing the kinetic of the wound width reduction in untreated (CT) or treated MCF10A (20 weeks, as indicated).

**A**

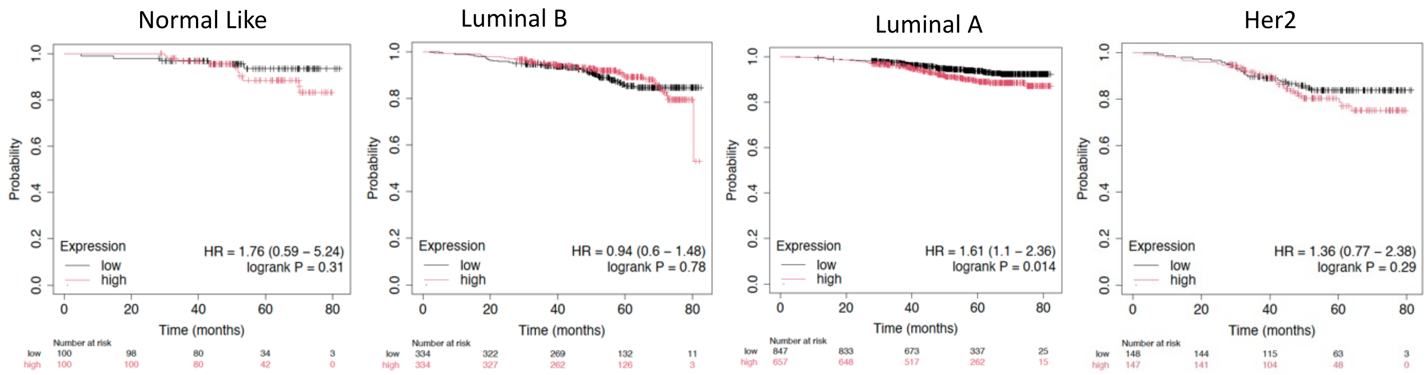


**B**

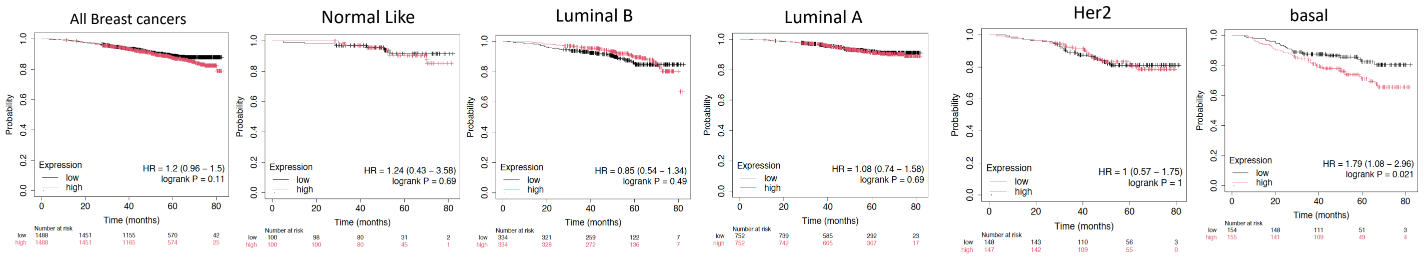


**Fig. S7. A.** Kaplan Meier analysis of the StemPlas signature in the indicated breast cancer subtypes**. B.** Kaplan Meier analysis of the 6 common genes up-regulated (HOXB6, KRT14, HSD17B2) or down-modulated (KHLH4, CNTNAP2 and RAB25) by 20 weeks of exposure to PS or PET in the indicated breast cancer subtypes**.**

**Table S1. Nanoplastic sources and physicochemical characteristics**

| | PS nanoplastic | PET nanoplastic |
|---|---|---|
| **Polymer type** | Polystyrene | Polyethylene terephtalate |
| **Source** | Commercial (Spherotech) | Real-life (water bottle milling) |
| **Particle shape** | Spherical | Irregular |
| **Particle size** | 46.65 ± 9.00 | 176.24 ± 6.52 |
| **Zeta average size** | 94.00 ± 0.86 water<br>189.63 ± 0.83 culture medium | 343.00 ± 18.90 water<br>402.43 ± 18.07 culture medium |
| **PDI** | 0.03 water<br>0.25 culture medium | 0.67 water<br>0.71 culture medium |
| **Zeta potential** | -44.90 ± 0.49 water<br>-12.7 ± 0.70 culture medium | -26.00 ± 0.84 water<br>-14.2 ± 0.05 culture medium |

**Table S2. List of Upregulated genes upon long-term exposure to plastic nanoparticles and included in the ENI10 signature.**

| PS and ENI10 | PET and ENI10 | ENI10-NP |
|---|---|---|
| | | AKR1B1 |
| AKR1B1 | FAP | AKR1B10 |
| AKR1B10 | | AKR1C1 |
| AKR1C1 | | C1S |
| C1S | | CD59 |
| CD59 | | CFB |
| CFB | | COL8A1 |
| COL8A1 | | CYP4B1 |
| CYP4B1 | | DCN |
| DCN | | FAP |
| FPR1 | | FPR1 |
| IL1RL1 | | IL1RL1 |
| LCN2 | | LCN2 |
| NAMPT | | NAMPT |
| NNMT | | NNMT |
| PAPSS2 | | PAPSS2 |
| PDZK1IP1 | | PDZK1IP1 |
| PI3 | | PI3 |
| PLSCR4 | | PLSCR4 |
| S100A7 | | S100A7 |
| SAA2 | | SAA2 |
| SAMSN1 | | SAMSN1 |
| SERPINA5 | | SERPINA5 |
| SERPINE2 | | SERPINE2 |
| SFTPB | | SFTPB |
| SLC39A8 | | SLC39A8 |
| SNCAIP | | SNCAIP |
| SRGN | | SRGN |
| SUSD2 | | SUSD2 |

**Table S3. Frequency of cells expressing surface markers quantified by flow cytometry after combined exposure to nanoplastic and BMP2.**

| | CD10+ / ECPAM- | CD44+ / CD24- | CD29+ / CD49f+ |
|---|---|---|---|
| **CT** | 8.57 ± 2.70 | 99.17 ± 0.30 | 97.55 ± 1.58 |
| **PS + BMP2** | 10.01± 3.00 | 98.90 ± 0.46 | 97.57 ± 1.04 |
| **PET + BMP2** | 11.33 ± 3.07 | 98.07 ± 0.70 | 98.08 ± 1.57 |
| **BMP2** | 11.02 ± 2.17 | 97.47 ± 0.70 | 98.98 ± 0.55 |
| **BMP2 + IL-6** | 15.07 ± 3.75 | 99.77 ± 0.10 | 98..85 ± 0.56 |
